# Pharmacodynamic encoding of tumor exposure identifies a mesenchymal state associated with antitumor sensitivity

**DOI:** 10.64898/2026.11.12.760322

**Authors:** Marta Rodriguez-Jiménez, Javier Robles, Ismael Fernández-Miranda, Raquel Altares, Maria José Guillen, August Vidal, Alberto Villanueva, Carmen Cuevas, Iñaki F Trocóniz, Marcelo Lima Ribeiro, Pablo Avilés

## Abstract

Predictive biomarker discovery in oncology relies on response phenotypes that do not account for tumor-site drug exposure. Here we developed PHENIX, a model-informed framework that uses tumor-exposure-driven pharmacodynamic parameters as quantitative phenotypes for molecular-state discovery. Using PM534, a colchicine-site tubulin inhibitor, we integrated measured drug concentrations with physiologically based pharmacokinetic and longitudinal pharmacokinetic–pharmacodynamic modeling. This yielded K_2_, quantifying the efficiency with which intratumoral exposure is converted into tumor damage across cell-derived xenograft models. Higher K_2_ was associated with a mesenchymal state enriched for epithelial-to-mesenchymal transition, focal adhesion and extracellular-matrix programs. An interpretable pathway-based model retained epithelial-to-mesenchymal transition and focal adhesion as candidate features and showed preliminary cross-dataset transferability. Proteomic analyses supported this molecular axis, whereas projection across CCLE and TCGA mapped its occurrence in contexts. Patient-derived orthotopic xenograft profiling provided orthogonal in vivo support. PHENIX demonstrates the feasibility of using model-derived pharmacodynamic parameters as molecular phenotypes for tumor-state discovery.

---

Predictive biomarker discovery in oncology commonly links baseline molecular features to nominal-dose response, terminal tumor volume, tumor growth inhibition, categorical responder status or in vitro viability. Although informative, these endpoints combine intrinsic growth, treatment schedule, tumor-site exposure and pharmacodynamic delay, making it difficult to distinguish insufficient tumor exposure from limited pharmacodynamic sensitivity^1^. Addressing this limitation requires response phenotypes that explicitly account for the exposure achieved at the tumor site.

Model-informed pharmacology provides a route to define more informative response phenotypes. Physiologically based pharmacokinetic (PBPK) models predict tissue exposure by integrating compound-specific properties with system-level physiology, whereas pharmacokinetic-pharmacodynamic (PK/PD) models link exposure to pharmacological effect.^2,3^ Because tumor uptake and intratumoral disposition can diverge substantially from plasma pharmacokinetics, intratumoral drug exposure is a critical but frequently underrepresented determinant of response interpretation.^4^

Nonclinical oncology studies typically evaluate antitumor activity in xenograft or patient-derived xenograft models using tumor growth inhibition, regression or survival as primary endpoints. Although these models remain indispensable for therapeutic evaluation, their conventional readouts are composite outcomes influenced by intrinsic growth kinetics, dosing schedule, intratumoral drug exposure and pharmacodynamic delay. Improving their translational value therefore requires moving beyond descriptive efficacy endpoints toward quantitatively interpretable exposure-response measures.^5,6^

When integrated with transcriptomic and proteomic profiling, xenograft models provide an opportunity to identify molecular states associated with quantitative pharmacodynamic response. Computational biomarker-discovery approaches increasingly link molecular features to drug sensitivity, but most still rely on in vitro viability, half-maximal inhibitory concentration, terminal response metrics or categorical sensitivity labels.^7^ Consequently, the molecular determinants of tumor response remain difficult to disentangle from drug exposure and treatment dynamics. Integrating a tumor-exposure-driven pharmacodynamic parameter with baseline molecular profiles could instead identify tumor states associated with the efficiency with which intratumoral exposure is converted into tumor damage.

To address this gap, we developed PHENIX (**Ph**armacodynamic **En**coding and Transcriptomic **I**ntegration for E**x**posure-Guided Oncology), a model-informed framework linking intratumoral drug exposure, longitudinal pharmacodynamic response and baseline molecular profiles. Using PM534, an optimized colchicine-site tubulin inhibitor, as a proof-of-concept system, we derived K_2_ as a quantitative measure of the efficiency with which intratumoral exposure is converted into tumor damage. Higher Kvalues were associated with a values were associated with a mesenchymal molecular state characterized by epithelial-to-mesenchymal transition (EMT) and focal adhesion programs. Pathway-based modeling, cross-dataset projection, proteomic profiling and patient-derived orthotopic xenograft (PDOX) pharmacodynamic studies provided complementary support for this association.

## RESULTS

### PHENIX integrates tumor exposure, longitudinal pharmacodynamics and molecular state

Conventional biomarker discovery typically associates baseline molecular features with terminal or categorical response outcomes, without explicitly accounting for tumor-site drug exposure, intrinsic growth kinetics or delay in pharmacodynamic response (Fig. 1a). To overcome these limitations, we developed PHENIX, a model-informed framework that integrates tumor pharmacokinetics, longitudinal pharmacodynamic response and baseline molecular profiling (Fig. 1b). PHENIX uses measured plasma, tissue and tumor drug concentrations to build a physiologically based pharmacokinetic model that reconstructs intratumoral drug exposure across dosing schedules (Fig. 1b,c). Predicted intratumoral drug concentrations are then linked to longitudinal tumor-volume dynamics through a semi-mechanistic pharmacokinetic–pharmacodynamic (PK/PD) model. This model yields K_2_, a quantitative parameter that captures the efficiency with which intratumoral drug exposure translates into tumor damage (Fig. 1d). Subsequently, K_2_ was integrated with baseline tumor molecular profiles to define a quantitative pharmacodynamic phenotype for transcriptomic, pathway-level, and machine-learning analyses (Fig. 1e).

**Fig. 1.**
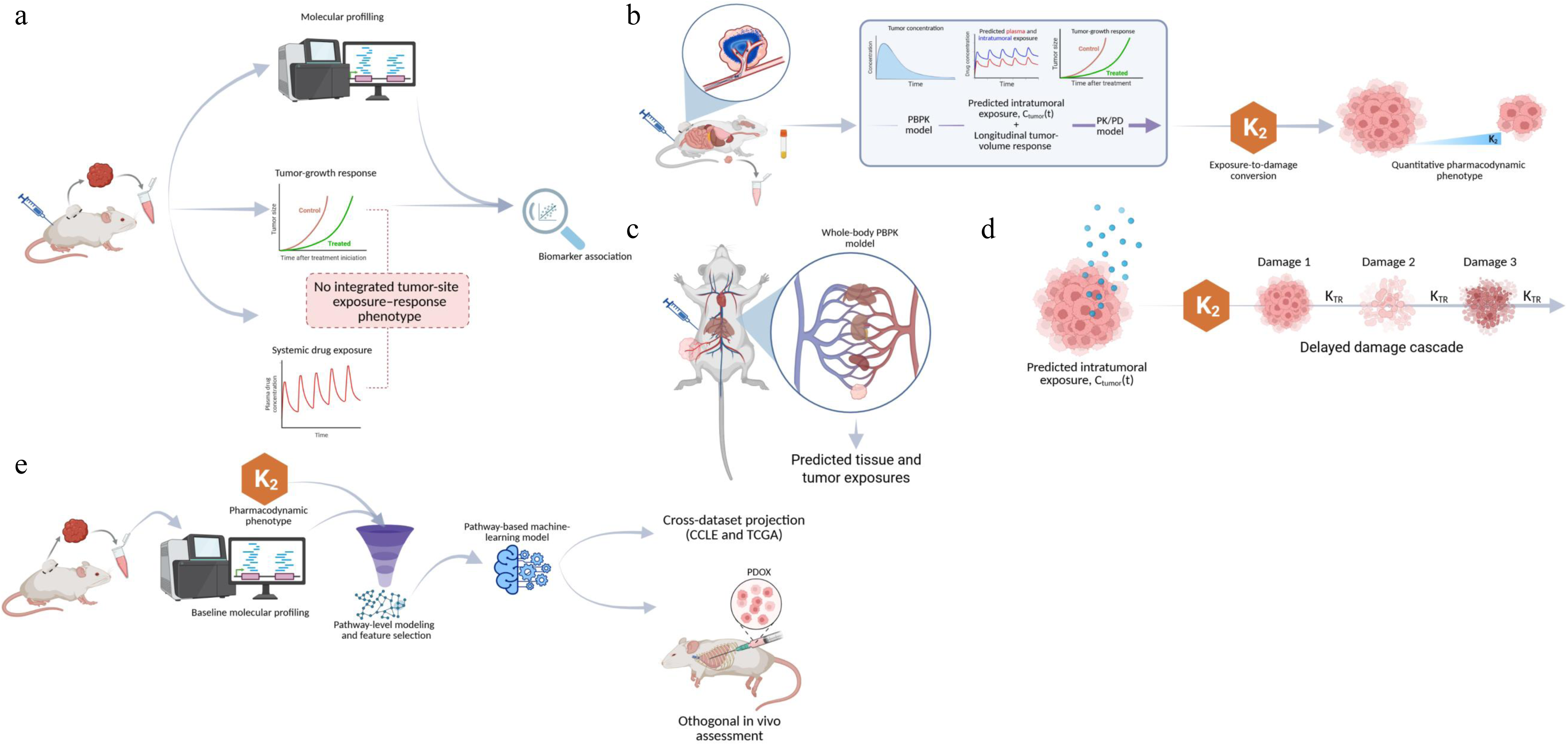
PHENIX integrates tumor exposure, pharmacodynamics and molecular-state discovery. (a) Conventional biomarker discovery links baseline molecular profiles directly to terminal or categorical tumor-response phenotypes, while systemic exposure and longitudinal tumor response are not explicitly integrated into a tumor-site exposure–response phenotype. (b) PHENIX integrates measured drug concentrations, PBPK-predicted intratumoral exposure and longitudinal tumor-volume response to derive K_2_ through tumor-exposure-driven PK/PD modeling. (c) Simplified whole-body PBPK architecture linking systemic circulation and representative tissue and tumor compartments to predict tissue and intratumoral drug exposure. (d) Tumor-exposure-driven PK/PD model in which predicted intratumoral exposure drives transfer of proliferating tumor volume into a delayed damage cascade through K_2_, with KTR governing transit through the damage compartments. (e) Molecular-state discovery workflow integrating baseline molecular profiles and the K_2_ pharmacodynamic phenotype through pathway activity scoring, feature selection and machine learning, followed by cross-dataset projection in CCLE and TCGA and orthogonal in vivo assessment in PDOX model.

Using PM534 as a proof-of-concept system, we applied PHENIX across CDX models with matched longitudinal tumor-volume data, PBPK–PK/PD-derived K2 estimates and baseline RNA-seq profiles (Fig. 1e). The CDX experimental design, engraftment conditions, dose levels and treatment schedules are summarized in Supplementary Table 1. This integrated design enabled us to test whether inter-model variation in pharmacodynamic sensitivity was associated with pretreatment molecular state and whether K2 captured molecular response features not adequately represented by nominal dose or terminal tumor-response endpoints.

### PBPK modeling reveals sustained intratumoral PM534 exposure

We first developed a PBPK model for PM534 using concentration-time data from plasma, tumors and major organs. Tumor concentration–time data from xenograft models were jointly used to characterize drug distribution within the tumor. The resulting tumor-exposure model was then used to predict intratumoral concentration–time profiles across the dose levels and treatment schedules evaluated across the broader CDX panel. Details of the bioanalytical LC-MS/MS (Liquid Chromatography–Tandem Mass Spectrometry) method used to quantify PM534 concentrations in plasma, tumors and organs are provided in Supplementary Table 2. The model reproduced the observed concentration-time profiles across sampled tissues and supported distinct pharmacokinetic behavior in tumors compared with non-tumor tissues. (Fig. 2a,b). The physiological characteristics used in the model and the estimates of the PBPK model parameters are listed in Supplementary Table 3. The full schematic and mathematical representation of the model is provided in Supplementary Material 1.

**Figure 2.**
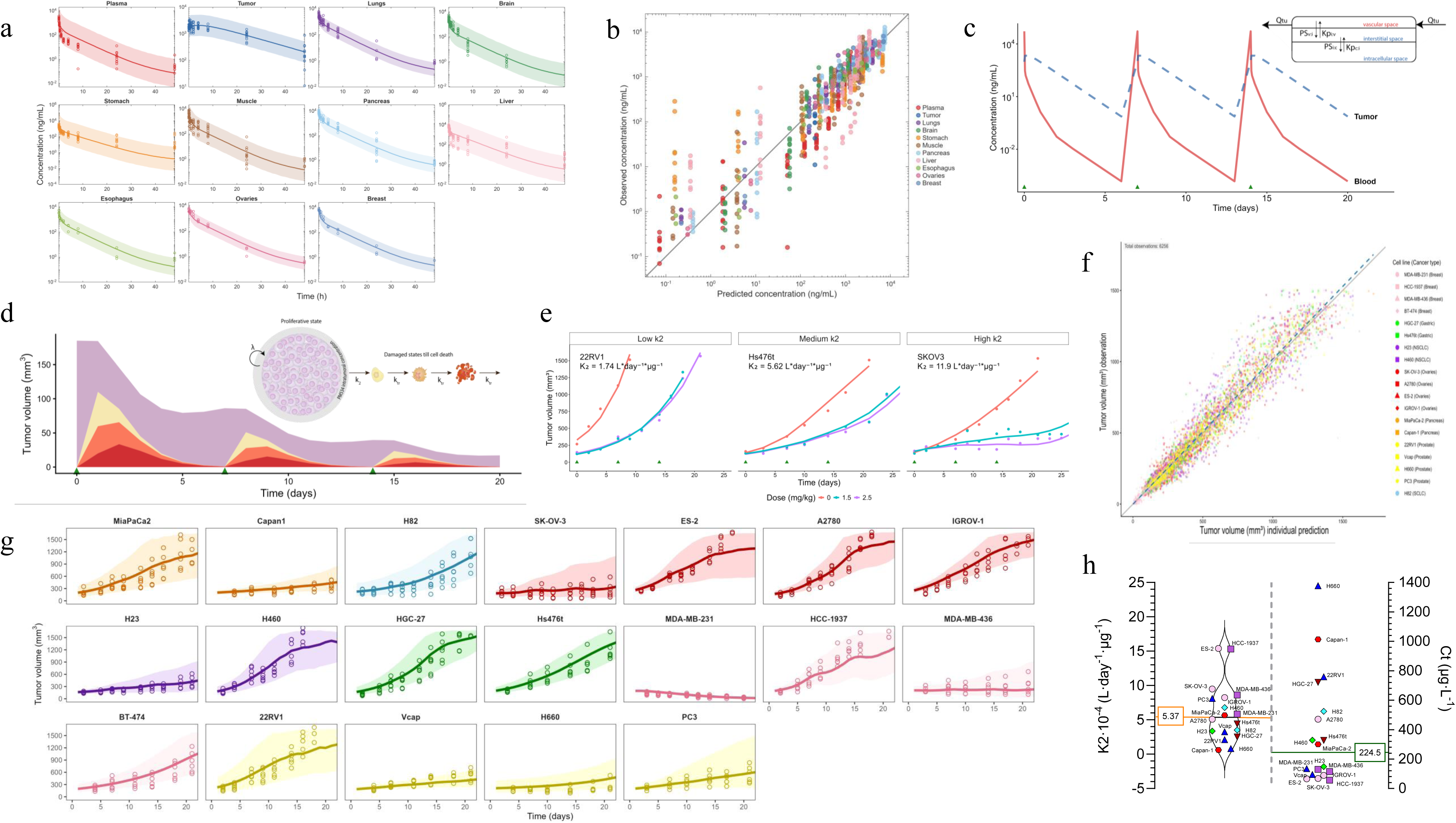
PM534 shows sustained intratumoral exposure and tumor-exposure-driven PK/PD modeling defines K_2_ pharmacodynamic sensitivity. (a) PBPK-predicted PM534 concentration-time profiles in plasma, tumor and selected tissues. Open circles show measured concentrations in different organs (raw data). Solid lines represent the median predicted concentrations, and shaded areas span the corresponding 95% prediction intervals; both were derived from 1,000 simulated studies. (b) Observed versus PBPK-predicted PM534 concentrations. (c) Schematic representation of the tumor compartment structure (vascular, interstitial and intracellular spaces) and PBPK-predicted intratumoral versus plasma PM534 exposure following 5 mg/kg intravenous administration on a q7dx3 schedule. Solid triangles indicate dosing time. (d) Model-predicted dynamics of tumor cell subpopulations following 5 mg/kg intravenous PM534 on a q7dx3 schedule. (e) Tumor volume vs time profiles following a q7dx3 dosing schedule for CDX models with low, intermediate and high estimates of K_2_. Different colors represent different dose levels. Solid lines represent model predictions. Solid circles, raw tumor volume data. (f) Observed versus model-predicted tumor volumes. (g) Observed and model-predicted tumor volume-time profiles for each CDX model under q7dx3 dosing, shown up to day 21 to allow comparison across models. Observations are represented as open points, while solid lines represent the mean of 1,000 simulations, and shaded areas the 90% prediction interval (5th–95th percentiles). (h) Distribution of model estimates of the K_2_ parameter (left) and the derived metric Ct (right). The values of 5.37 × 10 L·d ¹·µg ¹ and 224.5 µg/L are the medians across the 19 cell lines studied.

While drug distribution in most non-tumor tissues was adequately described by perfusion-limited model, tumor distribution required a permeability-limited model that distinguished between vascular, interstitial and intracellular compartments (Fig. 2c). PM534 concentrations in tumors were higher and more sustained than plasma concentrations, indicating that systemic pharmacokinetics alone did not fully represent target-site exposure (Fig. 2a). The parameter estimates were consistent with restricted intercompartmental exchange and prolonged tumor residence (Fig. 2c). Over the evaluated concentration-time interval, model-predicted tumor exposure exceeded plasma exposure, yielding a tumor-to-plasma AUC ratio of 35.

The fitted PBPK model was then used to predict intratumoral PM534 exposure under the dosing schedules used in the tumor growth inhibition experiments (Fig. 2c). These data provided the exposure basis for subsequent PK/PD modeling, in which predicted intratumoral PM534 concentrations, rather than nominal dose or plasma exposure alone, were used to drive tumor growth inhibition.

### Tumor-exposure-driven PK/PD modeling quantifies pharmacodynamic sensitivity across CDX models

Having established that PM534 exhibits sustained intratumoral exposure, we next investigated whether predicted tumor concentrations could quantitatively account for longitudinal tumor growth inhibition across CDX models. For each model, baseline tumor growth was first characterized using the most appropriate growth function in the absence of treatment and predicted intratumoral PM534 concentrations were then incorporated into a semi-mechanistic PK/PD model to describe drug-induced tumor damage (Fig. 2d).

Tumor growth and PK/PD parameter estimates for the CDX model are provided in Supplementary Table 4. The precision of the K_2_ parameters estimates, a key criterion for their use as a molecular phenotype in PHENIX, was assessed using relative standard errors (RSE). All K_2_ estimates met the predefined accepted threshold (RSE < 25%), except for those obtained in prostate tumor cell lines (PC3, 22RV1, and H660), which showed RSEs of 45-60%. The tumor-exposure-driven PK/PD model adequately captured treatment-induced tumor volume dynamics across CDX models and supported the use of a transit-compartment structure to account for the delay between intratumoral drug exposure and measurable tumor shrinkage (Fig. 2d). This modelling approach yielded K_2_ as tumor-exposure-driven pharmacodynamic parameter that quantifies the efficiency with which predicted intratumoral PM534 concentrations translate into tumor damage. The models also accurately described the observed tumor volume profiles and their variability (Fig. 2e–g).

K_2_ values varied substantially across CDX models, indicating marked heterogeneity in PM534 pharmacodynamic sensitivity despite use of the same compound and related dosing schedules (Fig. 2h). In addition, we also derived CT, calculated as λ/K_2_, as a model-derived estimate integrating intrinsic tumor growth and drug-induced damage into a predicted tumor concentration threshold associated with regression (Fig. 2h).

These analyses established K2 as the central quantitative response variable for downstream molecular modeling. Rather than representing terminal tumor size, categorical response or nominal-dose sensitivity, K2 quantifies the relationship between predicted intratumoral exposure and longitudinal tumor-damage kinetics.

### K_2_ captures a distinct dimension of antitumor response beyond conventional response metrics

To determine how K_2_ related to conventional antitumor response metrics, we performed principal component analysis integrating K_2_, terminal tumor-volume change (TTVC), tumor growth inhibition (TGI), best response, intrinsic tumor-growth rate (lambda) and CT (Fig. 3a; Supplementary Table 5). K_2_ contributed predominantly to PC1 (loading = 0.614), together with TTVC (loading = 0.562), whereas TGI and best response contributed mainly to PC2 (loadings = 0.683 and 0.708, respectively) (Fig. 3b). This distinction was further examined by cosine-similarity analysis. TGI and best response showed highly similar loading profiles in the PC1-PC2 space (cosine similarity = 0.993), whereas their similarities to K_2_ were substantially lower (0.157 and 0.036, respectively). By contrast, K_2_ and TTVC were closely aligned (cosine similarity = 0.996). Intrinsic tumor-growth rate made a comparatively limited contribution to the first two components. Together, these findings indicate that K_2_ captures a dimension of response distinct from TGI and categorical best response while remaining strongly related to terminal tumor-volume outcome.

**Fig. 3.**
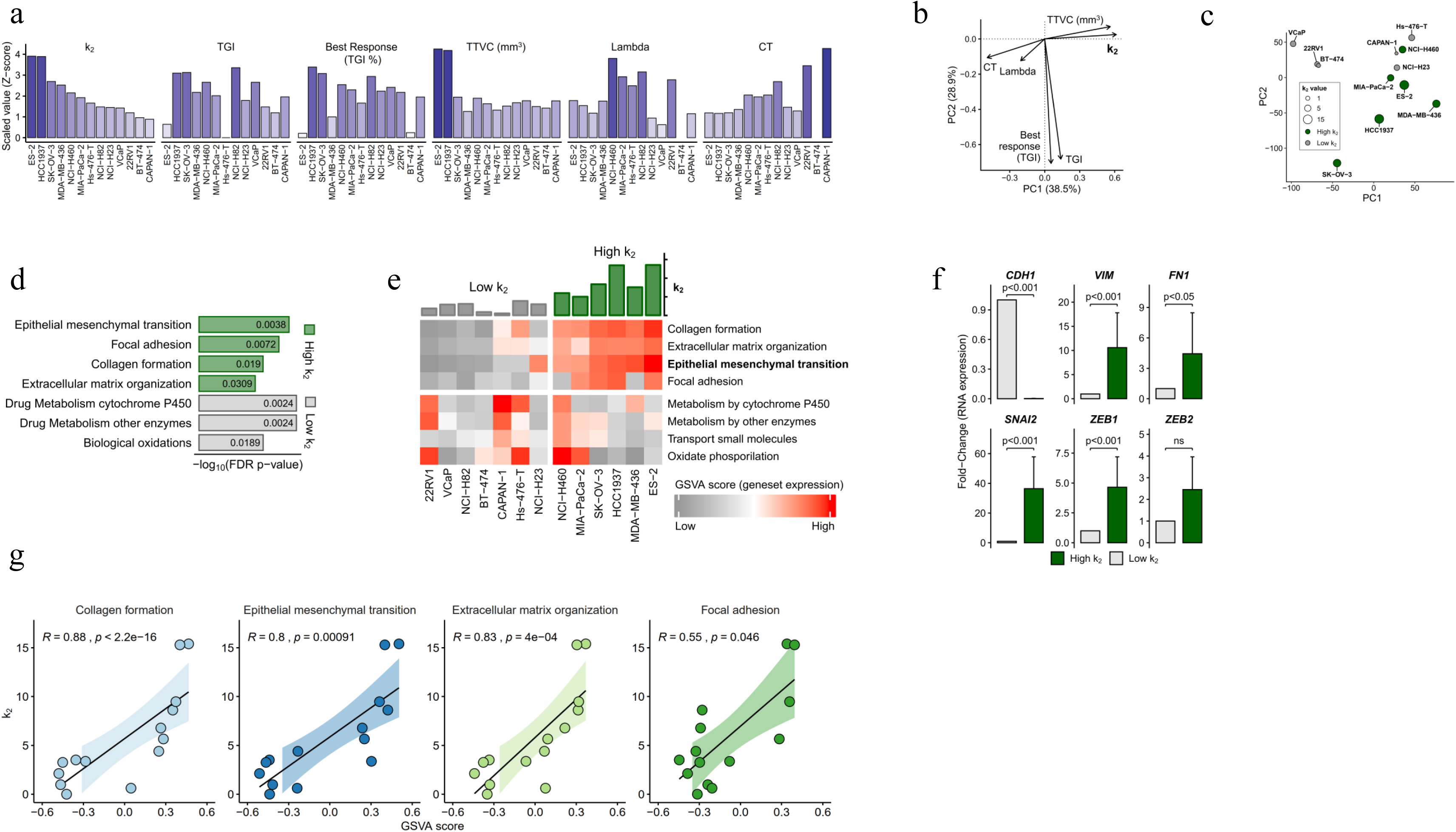
K_2_-defined pharmacodynamic sensitivity is associated with a mesenchymal EMT/focal-adhesion state. (a) Normalized distributions of K_2_ and conventional antitumor-response endpoints across CDX models, standardized to permit comparison among metrics measured on different scales. (b) Principal component analysis of K_2_ and antitumor-response endpoints. (c) Principal component analysis of baseline CDX transcriptomes, with models colored by K_2_ category. (d) Gene set enrichment analysis of epithelial-to-mesenchymal transition, focal adhesion, collagen formation and extracellular-matrix-related programs. The x axis represents −log10(FDR-adjusted P value). (e) Heatmap of representative genes involved in EMT, focal adhesion and extracellular-matrix organization. (f) Baseline expression of canonical epithelial and mesenchymal markers according to K_2_-defined sensitivity. (g) Associations between continuous K_2_ values and baseline pathway scores.

### K_2_ captures a mesenchymal and extracellular matrix-related transcriptional profile associated with PM534 sensitivity

To investigate whether baseline gene expression was associated with differences in PM534 pharmacodynamic sensitivity, we integrated pretreatment CDX RNA-seq profiles with corresponding PK/PD-derived K_2_ estimates. Tumor models were classified into high-and low-K_2_ groups using the cohort median for visualization, whereas continuous K_2_ values were used for pathway-level association analyses (Fig. 3c-g). RNA-seq processing metrics are provided in Supplementary Table 6.

Baseline transcriptomes showed a distinct distribution according to K_2_-defined sensitivity in the PCA space (Fig. 3c). High-K_2_ tumors were enriched for EMT, focal adhesion, extracellular matrix organization, collagen-containing matrix and migration-associated programs (Fig. 3d,e; Supplementary Table 7). Canonical mesenchymal genes, including VIM, FN1, SNAI2, ZEB1 and ZEB2, were increased in high-K_2_ tumors, whereas low-K_2_ tumors showed higher expression of epithelial and metabolism-associated programs, including CDH1-associated epithelial features (Fig. 3f).

Across pathway-level analyses, continuous K_2_ values were positively associated with baseline transcriptional programs related to extracellular matrix remodeling and mesenchymal features (Fig. 3g). The strongest association was observed for collagen formation (R = 0.88, P < 2.2 × 10□¹□), followed by extracellular matrix organization (R = 0.83, P = 4 × 10□□) and epithelial-mesenchymal transition (R = 0.80, P = 9.1 × 10□□) (Fig. 3g). Focal adhesion also showed a positive, although more moderate, association with K_2_ (R = 0.55, P = 0.046). Together, these results indicate that PM534 pharmacodynamic sensitivity is not randomly distributed across baseline molecular states but is preferentially associated with a coordinated mesenchymal and extracellular matrix-related transcriptional program involving collagen formation, matrix organization and cell adhesion.

### An interpretable pathway model retains EMT and focal adhesion as candidate features

The association between K_2_ and mesenchymal pathway activity prompted us to investigate whether baseline pathway states could predict pharmacodynamic sensitivity to PM534. To identify biologically relevant predictors, we first selected the 25 pathways most significantly associated with K_2_ based on gene set enrichment analysis and used their pathway activity scores as candidate predictors. We trained a sparse least absolute shrinkage and selection operator (LASSO) regression model using pathway activity scores as predictors and K_2_ as the response variable (Fig. 4a). The model retained two biologically coherent features, Hallmark EMT and KEGG focal adhesion, consistent with the differential expression, pathway enrichment and gene-level analyses (Fig. 4b). The candidate pathways and selected model coefficients are reported in Supplementary Table 8, and the corresponding K_2_ predictions are provided in Supplementary Table 9.

**Fig. 4.**
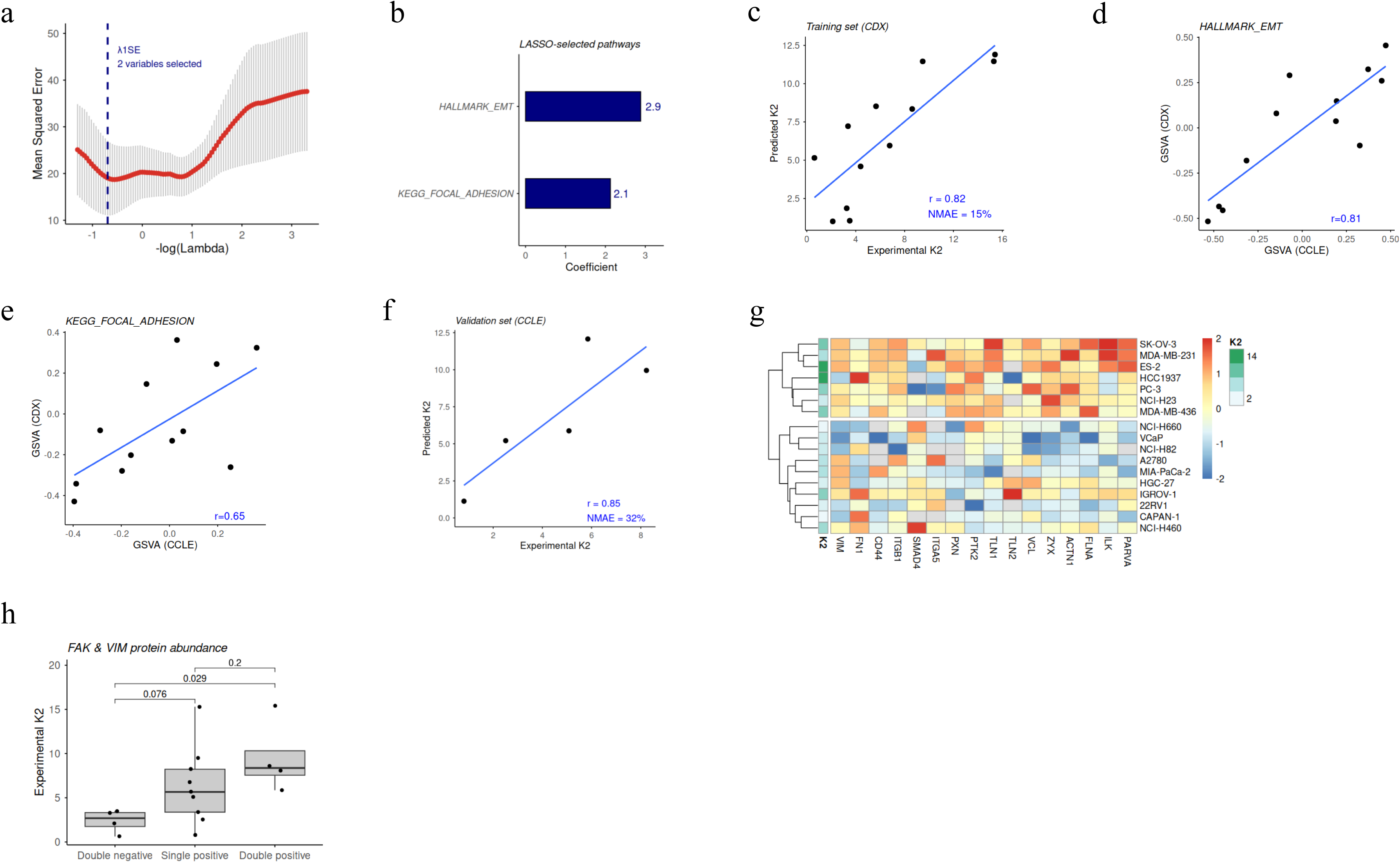
An interpretable pathway model retains EMT and focal adhesion as candidate features of PM534 pharmacodynamic sensitivity. (a) Ten-fold cross-validation curve used to select the optimal LASSO regularization parameter (λ). The λ.1se criterion was used to define the final sparse model. (b) Coefficients of the pathways retained in the final LASSO model. (c) Predicted versus experimentally estimated K_2_ values for the CDX training set. Pearson’s correlation coefficient (r) and normalized mean absolute error (NMAE) are indicated. (d, e) Concordance of Hallmark EMT (d) and KEGG focal adhesion (e) GSVA scores between CDX models and corresponding CCLE cell lines. Pearson correlation coefficients are shown. (f) Predicted versus experimentally estimated K_2_ values in the independent CCLE transfer set of five models. Individual models are labeled, and Pearson’s correlation coefficient (r), its 95% confidence interval and normalized mean absolute error (NMAE) are indicated (g) Heatmap showing the abundance of EMT-and focal adhesion-associated proteins across cancer models. Proteins detected in fewer than 70% of models were excluded. (h) Experimental K2 values stratified according to VIM and FAK protein abundance. Positive status was defined as protein abundance above the median value across all models.

In the CDX training set, predicted K_2_ values showed strong agreement with experimentally estimated K_2_ values (r = 0.82, NMAE = 15%), indicating that a model based on just two pathway activity scores, Hallmark EMT and KEGG focal adhesion, captured key features of PM534 pharmacodynamic sensitivity (Fig. 4c). To explore whether this model could be applied across datasets, we first evaluated the consistency of the two pathways activity in an independent cohort of CCLE-profiled cell lines with experimentally estimated K_2_ values. Both scores were correlated across datasets between datasets (EMT: r = 0.81; focal adhesion: r = 0.65), supporting the consistency of these transcriptional features across platforms (Fig. 4d,e). We then evaluated K_2_ predictions in an independent transfer set comprising five CCLE-profiled models. Predicted and experimentally estimated K_2_ values were strongly correlated (r = 0.85), although the prediction error was higher than in the CDX training set (NMAE = 31%; Fig. 4f). However, given the small sample size, the correlation estimate is subject to substantial uncertainty. These findings provide preliminary evidence of cross-dataset transferability rather than definitive external validation.

To assess the robustness of the pathway-based model, we evaluated feature-selection stability, out-of-sample predictive performance and prediction errors. Bootstrap resampling identified Hallmark EMT as the most consistently selected predictor, with KEGG focal adhesion also showing high selection frequency (Extended Data Fig. 1a; Supplementary Table 10). Leave-one-model-out cross-validation yielded moderate performance (r = 0.66, NMAE = 20%), indicating that predictions were not dominated by a single CDX model (Extended Data Fig. 1b). The observed prediction error lay near the lower tail of the permutation distribution, although the empirical test did not reach the conventional 0.05 threshold (P = 0.057; Extended Data Fig. 1c). Residual diagnostics showed no clear systematic bias across the prediction range (Extended Data Fig. 1d). Together, these findings indicate relative stability within the discovery dataset but warrant cautious interpretation given the modest sample size.

To explore whether the transcriptomic features identified by the pathway model were reflected at other molecular levels, we next examined proteomic profiles. Proteomic analysis provided orthogonal support for the EMT/focal adhesion axis identified by the transcriptomic pathway model. Expression patterns of EMT-and adhesion-associated proteins stratified cellular models according to K_2_-defined PM534 sensitivity, indicating that the biological signal captured by the model is reflected at the protein level (Fig. 4g). Notably, a descriptive two-marker representation based on VIM and FAK/PTK2 showed a directionally consistent relationship with K_2_-defined sensitivity (Fig. 4h), suggesting that the EMT/focal adhesion axis identified by the transcriptomic model is reflected at the protein level.

### PHENIX projects K_2_-associated molecular states across CCLE and TCGA

We next applied the PHENIX pathway model to large-scale transcriptomic datasets to determine whether the EMT/focal-adhesion state associated with higher K_2_ occurred across broader tumor contexts. In CCLE, PHENIX molecular-state scores were heterogeneous across cancer cell lines, indicating substantial variation in representation of the K_2_-associated transcriptional state (Fig. 5a). Lineage-level aggregation identified cancer contexts with higher PHENIX molecular-state scores, nominating them for prospective pharmacological evaluation. Model-derived scores and pathway activities across CCLE and TCGA are provided in Supplementary Tables 11 and 12.

**Fig. 5.**
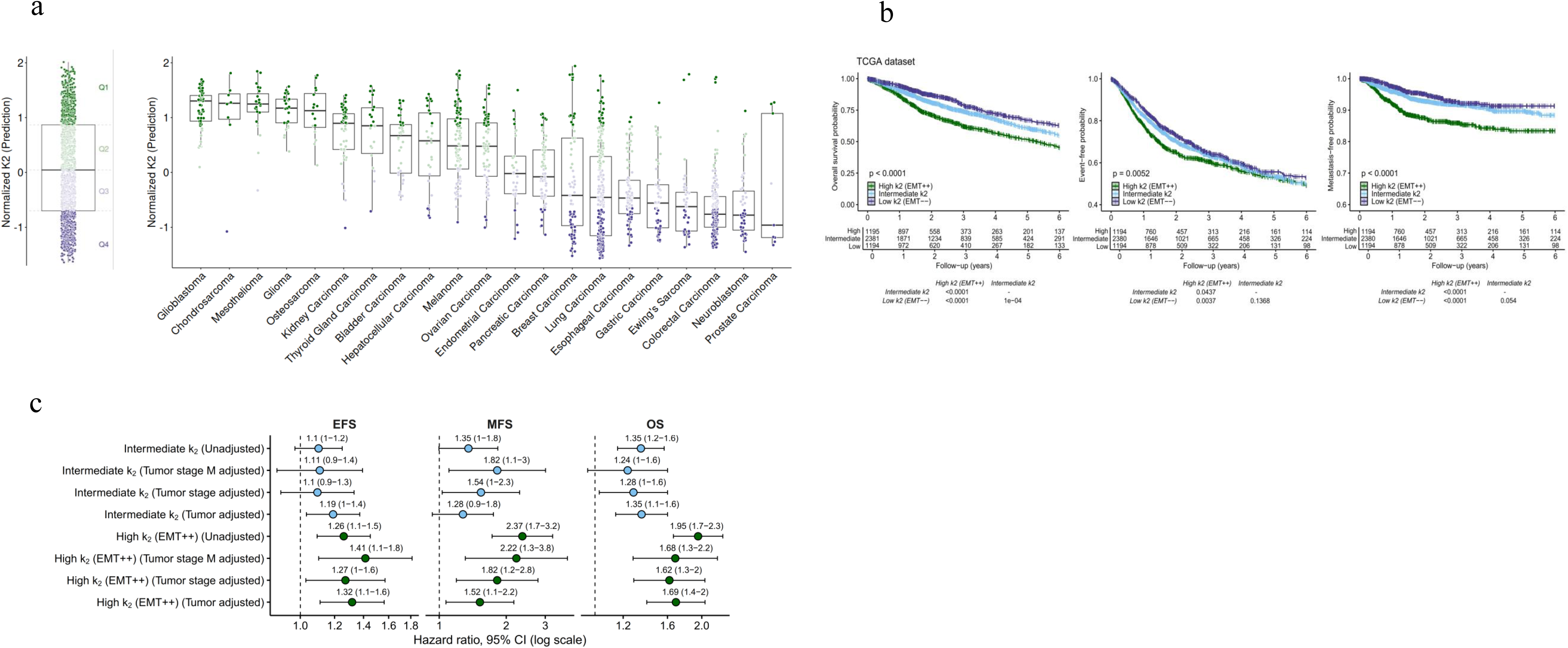
Projection of PHENIX K_2_-associated molecular states across CCLE and TCGA datasets. (a) Distribution of PHENIX molecular-state scores across CCLE cancer cell lines grouped by tumor lineage. Lineage-level aggregation nominates contexts for prospective pharmacological evaluation. (b) Human tumor-state mapping of the PHENIX molecular-state score across TCGA cohorts. Tumors were classified into high-, intermediate-and low-score groups, and associations with overall, event-free and metastasis-free survival were evaluated using Kaplan-Meier analyses. (c) Hazard ratios and 95% confidence intervals from univariable and multivariable Cox proportional-hazards models adjusted progressively for tumor type, pathological stage and pathological M status. These analyses characterize the prognostic context of the PHENIX molecular-state score and do not evaluate PM534 sensitivity or efficacy in patients.

We next projected the PHENIX pathway model onto TCGA tumors to determine whether the K_2_-associated molecular state occurred in human cancers and overlapped with clinically aggressive disease contexts. Because TCGA lacks PM534 treatment and tumor-exposure information, the model output was interpreted as a PHENIX molecular-state score rather than as a direct estimate of clinical drug sensitivity. Tumors with higher molecular-state scores were associated with poorer outcomes and aggressive disease features (Fig. 5b). Because EMT and focal-adhesion programs are themselves associated with aggressive disease biology, we used multivariable Cox proportional-hazards analyses to test whether these associations were maintained after accounting for tumor type, pathological stage and pathological M status (Fig. 5c). A high PHENIX molecular-state score remained associated with poorer overall, event-free and metastasis-free survival across the adjusted models. In the model accounting for tumor type and pathological stage, the high-score group had increased hazards of death (HR = 1.62, 95% CI 1.29-2.04; P < 0.001), disease progression or death (HR = 1.27, 95% CI 1.03-1.57; P = 0.026) and metastasis (HR = 1.82, 95% CI 1.19-2.78; P = 0.006). These associations were maintained after additional adjustment for pathological M status. Because TCGA does not include PM534 treatment or pharmacokinetic information, these analyses represent human tumor-state mapping and should not be interpreted as evidence of PM534 sensitivity or efficacy in patients.

### PDOX pharmacodynamic profiling provides orthogonal in vivo support

Because PHENIX identified EMT-and focal-adhesion-associated states as candidate PM534-sensitive contexts, we evaluated the transcriptional effects of PM534 in a clinically platinum-resistant ovarian cancer PDOX model. RNA-seq was performed on pretreatment, vehicle-treated and PM534-treated tumors (Fig. 6a). Relative to pretreatment tumors, vehicle-treated samples showed activation of EMT, focal adhesion, extracellular-matrix remodeling and migration-associated programs. PM534 treatment attenuated these changes relative to vehicle controls, reducing expression of a broad set of genes and suppressing migration-associated pathways (Fig. 6b,c). RNA-seq processing metrics are provided in Supplementary Table 6, and PDOX differential-expression outputs are provided in Supplementary Table 13.

**Fig. 6.**
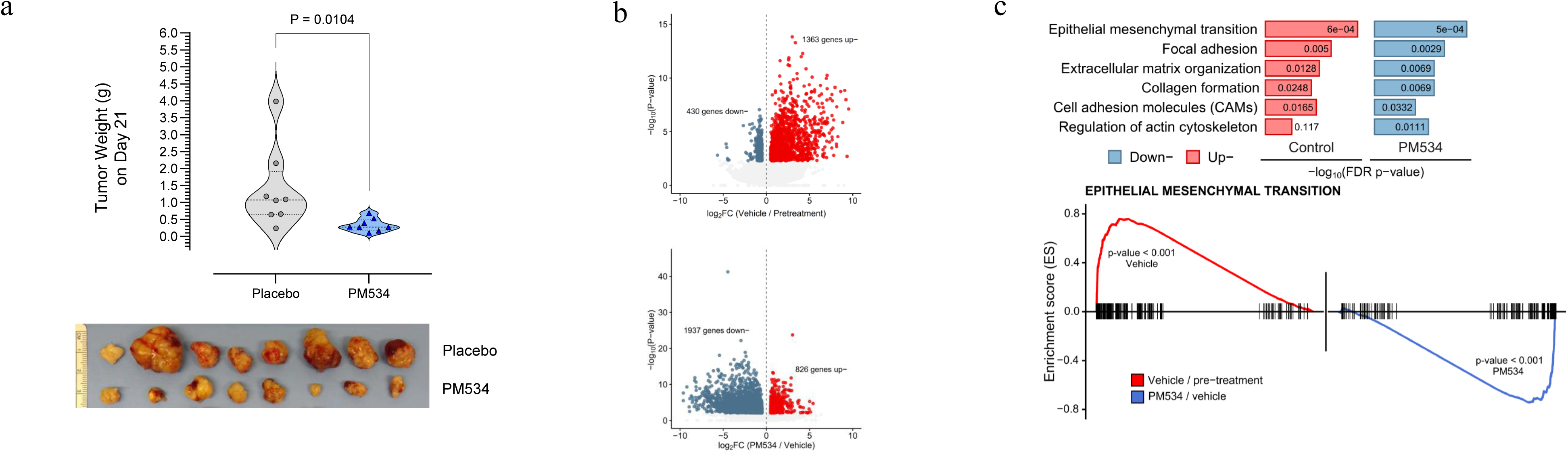
PDOX pharmacodynamic profiling provides orthogonal support for the PHENIX-defined tumor-state axis. (a) Upper panel: violin plots depicting tumor size in placebo-and PM534-treated mice at study completion (Day 21). Statistical significance was determined using Mann Whitney *U* test. Lower panel: representative gross photographs of excised tumors from each group (n=8) on Day 21. (b) Volcano plots showing differential gene expression in vehicle-treated versus pretreatment tumors and PM534-treated versus vehicle-treated tumors. Genes are plotted according to their log_2_ fold change and statistical significance, highlighting transcriptional changes associated with tumor progression and PM534 treatment, respectively. (c) GSEA of ranked gene lists derived from both differential expression comparisons, showing the pathway-level transcriptional changes associated with vehicle-treated versus pretreatment tumors and with PM534-treated versus vehicle-treated tumors.

These data do not constitute independent validation of the baseline predictor. Rather, they provide orthogonal pharmacodynamic evidence that PM534 treatment modulates the same EMT, focal-adhesion and extracellular-matrix programs associated with K_2_ in the discovery models.

## DISCUSSION

This study demonstrates the feasibility of using a model-derived tumor-exposure-driven pharmacodynamic parameter as a quantitative phenotype for molecular-state discovery. Rather than correlating baseline molecular features with nominal dose, terminal tumor volume, in vitro viability or categorical response, PHENIX derives K_2_ from PBPK-PK/PD modeling and asks which tumor states explain the efficiency with which intratumoral exposure is converted into tumor damage. The central advance is therefore not the characterization of PM534 alone, but the redefinition of the response phenotype used for biomarker discovery. In this framework, pharmacodynamics becomes more than a downstream measure of efficacy; it provides a quantitative bridge between intratumoral drug exposure, tumor biology and therapeutic response.

A key implication of this approach is that exposure should be considered at the tumor site. Administered dose or plasma pharmacokinetics may not adequately represent the concentration experienced by tumor cells when tumor penetration, retention, compartmental exchange or intracellular accumulation determines local exposure. Using plasma concentrations as the pharmacodynamic driver could still produce an adequate model fit, but K_2_ would then combine tumor distribution with pharmacodynamic activity. By anchoring the pharmacodynamic model to predicted intratumoral concentrations, PHENIX aims to reduce this confounding and provide a more biologically interpretable estimate of exposure-to-damage conversion. Tumor uptake has been highlighted as an underappreciated determinant of optimal dosing and translational response prediction.^2,3,4^

The use of K_2_ changed the biological question. Conventional response metrics, including terminal tumor volume, TGI and categorical responder status, reflect the combined influence of intrinsic growth, treatment duration, dose intensity, pharmacodynamic delay and sampling time. Tumor-growth inhibition models have long provided a quantitative framework for separating growth, drug effect and delayed tumor damage.^1^ PHENIX extends this principle by treating a drug-effect parameter as the phenotype to be explained by baseline molecular state. Accordingly, the transcriptomic and proteomic analyses go beyond identifying molecular features associated with observed outcomes to investigate which tumor states are associated with the efficiency with which intratumoral exposure is converted into tumor damage.

This distinction is mechanistically important. K_2_ captured a response dimension distinct from TGI and categorical best response, although it remained strongly related to terminal tumor-volume outcome. The value of K_2_ therefore lies not in being independent of antitumor efficacy, but in providing an exposure-anchored and longitudinally informed representation of response that can be interrogated molecularly.

Using this model-derived phenotype, PHENIX identified a mesenchymal EMT/focal adhesion state associated with higher PM534 pharmacodynamic sensitivity. This finding requires careful interpretation. EMT is classically linked to tumor plasticity, invasion, metastasis, stem-like features and therapeutic resistance^8^. However, EMT is not a uniform resistance state. Pan-cancer analyses have shown that EMT defines molecular states with distinct genomic, proteomic, immune and drug-response features^9^. In this context, our data suggest that EMT/focal adhesion activity marks a tumor state in which PM534 exposure is more efficiently converted into damage. The biological insight is therefore not that mesenchymal tumors are generally sensitive to PM534, but that an aggressive transcriptional state can encode a context-dependent pharmacological vulnerability.

This interpretation is plausible for a compound targeting tubulin-dependent cytoskeletal organization, but it remains mechanistically unproven. EMT and focal adhesion programs are linked to cytoskeletal remodeling, cell-matrix interaction, mechanical signaling and migration-associated phenotypes, any of which may influence dependence on microtubule dynamics or adhesion-associated survival programs^10–12^. The convergence of differential expression, pathway enrichment, continuous K_2_-pathway association, sparse pathway modeling and proteomic support argues that the EMT/focal adhesion axis is not an arbitrary correlation. However, perturbation experiments targeting EMT regulators, focal adhesion signaling or cytoskeletal-state dependencies will be required to determine whether this state is causal, permissive or a composite marker of PM534 vulnerability.

The pathway-based predictor was intentionally designed for interpretability rather than black-box performance. Sparse LASSO modeling selected EMT and focal adhesion, two biologically coherent features that aligned with independent transcriptomic and proteomic analyses. This is appropriate for biomarker discovery, where feature interpretability is as important as predictive accuracy, but small-sample, high-dimensional models remain vulnerable to instability and overfitting.^13^ Accordingly, the current PHENIX predictor should be viewed as a tumor-state prioritization tool, not as a calibrated clinical classifier. The increased absolute prediction error observed during cross-dataset transfer is consistent with recognized limitations of machine-learning-based cancer drug-response prediction, including differences in sample origin, normalization, platform scale and biological context.^14^

The external analyses provide a hierarchy of support rather than equivalent validation. The five-model CCLE transfer set provided preliminary evidence of cross-dataset transferability. Projection across the broader CCLE collection nominated molecularly defined cancer contexts for prospective pharmacological evaluation. TCGA established that the same molecular-state score occurs in human tumors and overlaps with aggressive disease biology but did not evaluate PM534 sensitivity or efficacy. PDOX profiling provided orthogonal pharmacodynamic evidence that PM534 treatment modulates EMT, focal-adhesion and extracellular-matrix programs in a clinically resistant in vivo model.

PHENIX complements model-informed drug development, which increasingly uses mechanistic PK/PD and exposure–response analyses to support oncology dose and schedule optimization. ^15^ The FDA Oncology Center of Excellence’s Project Optimus reinforces this shift from maximum-tolerated-dose selection toward integration of efficacy, safety, tolerability and exposure–response evidence.^15–18^ Whereas these approaches primarily inform which regimen should advance, PHENIX asks which tumor molecular states most efficiently convert intratumoral exposure into pharmacodynamic response. It may therefore help prioritize tumor contexts for subsequent dose–schedule evaluation but does not replace clinical dose-finding or optimization.

By integrating longitudinal tumor growth, intratumoral exposure and molecular profiling, PHENIX extracts more information from each in vivo experiment than terminal-response comparisons alone. The framework may help prioritize informative dose–schedule studies and focus confirmatory work on tumor states with a strong exposure–response rationale, consistent with the 3Rs principle of refining animal use. ^19,20^ PHENIX is not intended to replace animal studies, but to support their more selective design and quantitative interpretation.

This study has limitations. PHENIX was developed using one compound, and extension to additional mechanisms of action will be required to establish broader generalizability. K_2_ estimates depend on the adequacy of the PBPK–PK/PD model, including assumptions about tumor exposure, compartmental exchange, delayed drug effect and model identifiability. In addition, direct tumor concentration–time data were available for four xenograft models, whereas intratumoral exposure in the broader CDX panel was reconstructed using the resulting tumor-distribution model. Because model-specific tumor perfusion, vascular fractions and permeability were not measured across the complete panel, the analysis assumes that the PBPK structure adequately captures the principal determinants of tumor disposition across models. Model-specific differences in tumor vascularity, stromal composition or permeability could therefore contribute to uncertainty in the reconstructed intratumoral exposures and, consequently, in the estimated K_2_ values. The predictive model was trained on a modest number of CDX models, making feature stability and overfitting important concerns. Cross-dataset transfer between CDX, CCLE, TCGA, PDOX introduces differences in sample origin, stromal content, normalization and platform scale.

More broadly, PHENIX illustrates how pharmacological response can be modeled as a quantitative molecular phenotype. Recent neural-PK/PD and machine-learning approaches highlight the convergence between pharmacometric modeling and computational prediction of longitudinal drug-response dynamics.^21^ PHENIX takes a complementary approach, rather than replacing mechanistic PK/PD modeling with a black-box predictor, it uses a mechanistically derived PK/PD parameter as the phenotype to be explained by molecular data. PM534 served as the experimental system through which this strategy was developed and tested, but the conceptual opportunity is broader. By connecting tumor exposure, pharmacodynamic damage and baseline molecular state, PHENIX provides a proof-of-concept framework for identifying tumor contexts in which pharmacological exposure is most efficiently converted into tumor control.

PHENIX provides a proof of concept for using tumor-exposure-driven pharmacodynamic parameters as quantitative phenotypes for molecular-state discovery. In the PM534 system, efficient exposure-to-damage conversion was associated with a mesenchymal EMT/focal-adhesion state. Additional compounds, larger independent cohorts and mechanistic perturbation studies will be required to establish generalizability and predictive utility.

## ONLINE METHODS

### Pharmacokinetic/Pharmacodynamic modeling

Model development was performed sequentially. A pharmacokinetic model was developed first, followed by characterization of tumor growth under unperturbed conditions and by a pharmacodynamic model describing PM534-induced antitumor response. This sequential strategy provided the quantitative pharmacodynamic phenotype used by PHENIX to link tumor exposure, longitudinal response and baseline molecular state.

### Pharmacokinetic Modeling

A physiologically based pharmacokinetic (PBPK) model was developed in mice using drug concentrations measured in blood, tumor and several additional organs. The model structure is represented schematically in Fig. 1c, and the tumor vascular, interstitial and intracellular subcompartment structure is represented in Fig. 2c and Supplementary Material 1, for a full schematic and mathematical description of the PBPK model.

Under the assumption of perfusion-limited distribution, each organ is fully characterized by a single homogeneous compartment, with drug input and output rates equal to with drug input and output rates equal to φ_Org ×_ C_Arterial_ and 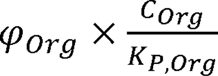, respectively, where C_Arterial_ represents the drug concentrations in arterial blood, and φ_Org_, C_Org_, and K_P,Org_ refer to the blood flow, tissue drug concentration, and partition coefficient between tissue and blood for each specific organ, respectively. Based on the known pharmacokinetic characteristics of PM534, drug clearance (CL) was assumed to occur by liver metabolism. The values of cardiac output, organ-specific blood flows and organs volumes used in this analysis were taken from literature ^22,23^ or calculated according the weight of the mice using algorithms implemented in PK-sim® (Open Systems Pharmacology Suite 12) and are listed in Supplementary Table 3. For the tumor, total volume was extrapolated from experimental measurements, while tumor-blood flow and its vascular and interstitial volume fractions were obtained from literature.^24–26^

For permeability-limited distribution, by contrast, each compartment is further divided into three sub-compartments (vascular, interstitial and intracellular) and, in addition to K_P,Org_, and CL, parameters accounting for permeability are estimated.

Several questions arose during the analysis and were addressed during model building. The main ones were: (i) whether the pharmacokinetics of PM534 differed between tumor-bearing and non-tumor-bearing mice; (ii) whether drug uptake into the tumor differed across cell lines; and (iii) whether organ distribution of the drug was perfusion-or permeability-limited.

The pharmacokinetic analysis was conducted following a naïve pooled-data approach, since each animal contributed a single observation for each of the organs studied. Residual error, defined as the difference between the observed concentrations and the model predictions, was characterized by an additive model on a logarithmic scale, and its magnitude was estimated separately for each organ.

### Tumor/plasma exposure ratio calculation

Tumor-to-plasma exposure ratios were calculated from model-predicted concentration-time profiles (C_pred_) to quantify divergence between systemic and intratumoral PM534 exposure.

The area under the concentration-time curve from time 0 to the last measured time point (AUC_0-tlast_) was calculated using the following expression which integrate either plasma or tumor exposure over time:

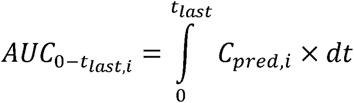

Where *i*, refers to either plasma or tumor concentrations. Tumor/plasma exposure ratio (AUC_Ratio_) was calculated as: AUC_Ratio_ = AUC_0-tlast,tumor_ / AUC_0-tlast,plasma_.

### Disease and pharmacokinetic/pharmacodynamic models

The longitudinal tumor volume data from each cell line were analyzed separately. Tumor volume measurements were logarithmically transformed during the analysis.

The ability of the exponential, Gompertz, and so-called Simeoni^1^ models to describe tumor growth dynamics in the groups of animals receiving saline solution (control groups) was evaluated. In the aforementioned models, the contribution of exponential growth is pivotal: Equation 1 describes the exponential rate of change of tumor volume (TV) in the absence of pharmacological perturbation and represents the core structure into which drug effects are embedded, as will be shown below.

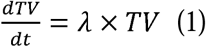

Where λ is the first-order rate constant of tumor growth and the system is initialized at TV, the initial tumor volume. It is worth noting that λ lumps together the (im)balance between tumor cell progression and the natural death of tumor cells; however, these individual contributions could not be separately identified.

Tumor shrinkage is driven by the predicted drug concentrations in the tumor (C_Tumor_). The selected PBPK model, with its corresponding parameters, was used to generate drug concentrations following the dosing schedules (a dose every week for 3 consecutive weeks; q7dx3) used in the tumor growth inhibition experiments. Figure 2c shows the PBPK model-based predictions obtained in plasma and tumor after (q7dx3) administration of a 5 mg/kg intravenous dose. Drug effects were incorporated in the model as follows (equation 2):

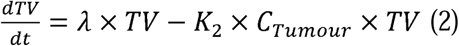

Where K_2_ is a second-order rate constant governing drug-induced tumor cell death.^1,27^ The K_2_ parameter allows different drugs to be ranked for a given tumor type, or the sensitivity to a specific drug to be ranked across different tumor types. The greater the estimate of K_2_ the higher tumor shrinkage. The model represented by equation 2 implies a direct and instantaneous effect with respect to C_Tumor_ levels. Although such a rapid onset of response can be observed in other therapeutic areas (e.g., cardiovascular), in oncology anti-cancer drugs typically exert their effects through intracellular conformational changes that require a certain period to appear and that cause the effects to persist for some time after drug levels have declined due to elimination. This delay in the onset and offset of tumor-shrinkage effects is incorporated into our model framework through a series of three transit compartments, each representing tumor cells (TV_Damage1-3_) with a different degree of damage severity (equations 3-5):

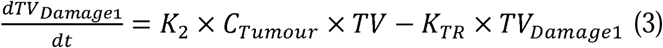

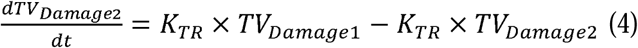

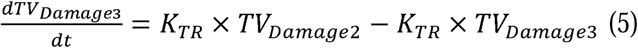

Where K_TR_ is the first order rate constant that governs the delay in the onset and offset of drug effects. The initial conditions of TV_Damage1-3_ are equal to 0. Figures 2d show the time course of TV_Damage1-3_ and TV, respectively, corresponding to different values of K_TR_. Higher values of K_TR_ are associated with shorter onset and offset response times. Total tumor volume is the results of the sum of the different volumes (TV + TV_Damage1-3_).

Different model structures were also considered during the model-building process: (i) resistance development, (ii) clone heterogeneity with respect to drug sensitivity, and (iii) saturation of drug effects.

The pharmacokinetic analysis was conducted following the Non-linear Mixed Effects Approach. Inter-animal variability in model parameters was modelled exponentially, and the residual error was described with an additive model on logarithmic scale.

### Conventional response metrics and endpoint comparison

To evaluate the added value of K_2_ relative to conventional antitumor response metrics, endpoint values were calculated at the tumor-model level. Terminal tumor volume change (TTVC) was calculated as:

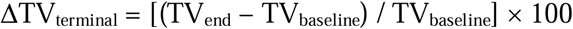

Tumor growth inhibition was calculated as:

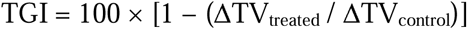

where:

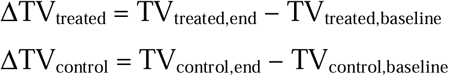

Best response was calculated as the minimum percentage change from baseline tumor volume observed during treatment:

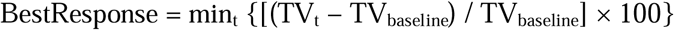

A principal component analysis (PCA) was performed to compare K_2_ with conventional and model-derived antitumor response metrics across CDX models. The analysis included K_2_, TTVC, TGI, best response, intrinsic tumor growth rate (λ), and CT. All variables were converted to numeric format, and metrics with insufficient observations or no variance were excluded. Missing values were imputed using the median of the corresponding metric. Because the included endpoints were measured on different scales, variables were centered and scaled to unit variance before PCA. The contribution of each response metric to the first two principal components was assessed from its PCA loadings. Similarity between metrics was additionally quantified as the cosine similarity between their loading vectors in the PC1– PC2 space, with values approaching 1 indicating aligned loading profiles, values approaching −1 indicating opposite orientations, and values close to 0 indicating approximately orthogonal profiles. PCA scores were used to visualize the distribution of CDX models, whereas loading plots were used to assess the relationships among response metrics. The CDX endpoint values included in the PCA and the resulting loading-based statistics are reported in Supplementary Table 5.

### Model Selection and Evaluation

Selection between competing model candidates was performed attending to the (i) value of the Akaike Information Criteria,^28^ (ii) visual inspection of the goodness of fit plots, and (iii) parameter identifiability reflected in the magnitude of the relative standard errors [RSE(%)] calculated as the ratio (multiplied by 100) between the standard error and the point estimate of the parameter.

Model performance was evaluated through simulation based diagnostics, where 1000 studies of the same characteristics of the original ones were simulated using the selected models and their parameter estimates. For each virtual study and observation time, the median of the simulated values was calculated and then, the 2.5^th^ and 97.5^th^ percentiles of the median vales were represented together with the raw data.

### Software

Data visualization and model evaluation were performed using R version 4.3.0 through RStudio interface. Population PK-PD analyses were conducted using NONMEM® version 7.5.1 (ICON PLC, Ireland).

### Cell-Derived Xenograft Tumor Models

All animal protocols were reviewed and approved according to regional Institutional Animal Care and Use Committees (PROEX 121.3/21 and 212.8/23).

Athymic Nude-Foxn1*^nu/nu^* and CB17/Scid mice were supplied by Envigo RMS (USA). Mice were housed (five per cage) in individually ventilated cages (Sealsafe^®^ Plus, Techniplast S.P.A.; Italy), on a 12-hour light-dark cycle at 21-23 °C and 40-60% humidity. Mice were allowed free access to irradiated standard rodent diet (Tecklad 2914C. Envigo RMS; USA) and sterilized water. Animals were acclimated for five days prior to being individually tattoo-identified.

Design, randomization and monitoring of experiments (including body weights and tumor measurements) were performed through NewLab software v2.50.1.6 (NewLab Oncology; France).

All tumors were engrafted in female athymic Nude-Foxn1*^nu/nu^*mice except VCap tumor (prostate) or BT-474 (breast) that was implanted in male or female CB17/Scid mice respectively. A summary of experimental details is provided in Supplementary Table 1.

When tumors reached 150-200 mm^3^, mice (n≥6 mice/group) were randomly allocated (Day 0) to receive PM534 or placebo. PM534 doses were selected based on the previously calculated maximum tolerated dose (MTD), that was determined to be 4.75 mg/kg (2.5 mg/kg in male CB17/Scid mice).^29^ Treatment was weekly intravenous administration for two or three consecutive weeks (days 0, 7 and 14). Caliper measurements of the tumor diameters were performed 2 to 3 times a week, and tumor volumes were calculated according to the following formula: volume =(a·b^2^)/2, where *a* and *b* presented the longest and shortest tumor diameters, respectively. For survival evaluation, the euthanasia endpoint was either a tumor volume larger than 2,000 mm^3^, tumor necrosis, or both.

### Generation and characterization of a cisplatin-resistant ovarian cancer patient-derived orthotopic xenograft (PDOX) model and evaluation of its response to PM534 treatment

The primary tumor specimen was obtained at Hospital Universitari de Bellvitge (L’Hospitalet de Llobregat, Barcelona, Spain). The study was approved by the institutional review board, and written informed consent was obtained from the patient, who had not previously received cisplatin-based chemotherapy. PDOX experiments were conducted at the Parc Científic de Barcelona (PCB) in accordance with the procedures approved by the Ethical Committee of Animal Experimentation of the Parc Científic de Barcelona and Catalan Government (23-049-P4) and complied with the relevant ethical guidelines for the care and use of laboratory animals.

Non-necrotic fragments (approximately 2–3 mm³) from a resected human high-grade serous ovarian carcinoma (HGSOC) were maintained in DMEM (BioWhittaker) supplemented with 10% fetal bovine serum (FBS) and penicillin–streptomycin at room temperature until implantation. Under isoflurane anesthesia, mice underwent a lateral laparotomy to expose the ovaries, and tumor fragments were sutured to the ovarian surface using 7-0 Prolene sutures. Tumor growth was monitored two to three times per week. Once established, tumors were harvested, divided into fragments, and serially transplanted into two to five recipient mice. During early passages, tumor fragments (6–8 mm³) were cryopreserved in liquid nitrogen in 90% FBS and 10% dimethyl sulfoxide (DMSO) for subsequent implantation.

The cisplatin-resistant derivative (R-HGSOC) was established through repeated cycles of in vivo cisplatin treatment and tumor relapse, as previously described^30^ (data not shown). Briefly, orthotopically engrafted HGSOC tumors at mouse passage 3 were allowed to grow until palpable intra-abdominal masses developed. Tumor-bearing mice received intravenous cisplatin once weekly on days 0, 7 and 14, starting at 2 mg/kg. The dose was progressively increased across successive treatment cycles to 3, 3.5, 4 and 5 mg/kg. After tumor relapse, tumors were harvested and reimplanted into new recipient mice for subsequent treatment cycles.

The morphology of the primary tumor and the HGSOC and R-HGSOC xenografts was compared by hematoxylin and eosin (H&E) staining of paraffin-embedded sections. Immunohistochemical analyses of cytokeratin 7 (CK7), Ki-67, WT1, estrogen receptor (ER) and progesterone receptor (PR) were performed according to the standard clinical protocols of the Department of Pathology (data not shown).

To evaluate the response to PM534, female mice were orthotopically implanted with R-HGSOC tumor fragments. Once tumors reached a comparable palpable size, animals were randomly allocated to the placebo or PM534 treatment group (n = 8 per group). PM534 was administered intravenously at 4.75 mg/kg once weekly for three consecutive weeks (days 0, 7 and 14). Placebo-treated animals served as controls. On day 21, seven days after the final treatment, mice were euthanized, and the ovaries were dissected and weighed. Representative tumor fragments were either snap-frozen in liquid nitrogen or fixed and processed for paraffin embedding. Humane endpoints were defined as body weight loss exceeding 20% for more than 48 h or the development of signs of pain, distress or severe morbidity. PDOX samples used for transcriptomic pharmacodynamic analyses are described in the PDOX transcriptomic pharmacodynamic analysis section.

### Pharmacokinetics and distribution studies in mice

Animals bearing tumors (150-200 mm^3^) received a single bolus intravenous dose (4.75 mg/kg) of PM534. Then, blood, tumors (A2780 -ovary-, HGC-27 -gastric-, H460 -NSCLC-and MDA-MB-231 -triple negative breast-) and organs (brain, breast, liver, lung, ovary, pancreas, skeletal muscle -*rectus femoralis* and *vastus medius*, *lateralis* and *intermedius*-and stomach) were collected at 0.083 (5 min), 0.25 (15 min), 0.5 (30 min), 1, 2, 4, 8, 24 and 48 hours post-dose (n=3 mice/sampling time) from CO_2_ euthanized animals.

The blood was obtained by cardiac puncture and transferred to tubes containing K_3_ EDTA as an anticoagulant, and plasma was then collected after centrifugation at 3,000 rpm for 15 minutes at 4 °C. Once dissected free, tumors and organs were weighed and then diluted 1:4 (w/v) with ammonium hydroxide 0.5 N and homogenized for 20 seconds at 6,800 rpm twice (Precellys Evolution, Bertin Instrument, France). The resulting plasma, tumor and organ extracts were divided into aliquots and frozen (−80°C) until bioanalysis, which consisted of supported liquid-liquid extraction (SLE), followed by gradient reverse-phase chromatography and detection by positive-ion electrospray tandem mass spectrometry (ESI/MS/MS) using multiple reaction monitoring (MRM). Bioanalytical method details are provided in Supplementary Table 2.

Non-compartmental pharmacokinetic analysis of PM534 was performed using Phoenix^®^ WinNonlin^®^ version 6.3 (Certara L.P. (Pharsight); USA) according to a composite sampling method.

### RNA extraction, library preparation, and RNA sequencing

Total RNA was isolated using the RNeasy Mini Kit (Qiagen, Cat. #74106), following manufacturer’s instructions. RNA concentration and integrity were evaluated using a NanoDrop spectrophotometer (Thermo Fisher Scientific) and an Agilent 2100 Bioanalyzer (Agilent Technologies). Samples with RNA Integrity Number (RIN) ≥ 8 were selected for downstream sequencing. RNA-seq libraries were prepared using the Illumina TruSeq Stranded mRNA Library Prep Kit (Illumina, Cat.# 20020595). Briefly, mRNA was enriched using oligo(dT) magnetic beads, fragmented, and reverse transcribed into cDNA. The resulting cDNA was end-repaired, ligated to Illumina adapters, and PCR-amplified. Library quality and fragment size distribution were verified using an Agilent Bioanalyzer. Sequencing was performed on an Illumina NovaSeq 6000 platform to generate at least 30 million paired-end 100 bp reads per sample.

### RNA-seq bioinformatic analysis

Raw sequencing data were processed using the nf-core/rnaseq pipeline (latest stable release), ensuring a fully reproducible and standardized workflow. Raw reads were assessed using FastQC and adapters were trimmed with Trim Galore. Given that the RNA-seq data were derived from CDX models, reads were aligned in parallel to both the human reference genome (GRCh38) and the mouse reference genome (GRCm39) to account for potential host (murine) contamination. Alignment was performed using STAR, a splice-aware aligner, against each reference genome independently. Gene-level counts were quantified using featureCounts based on the corresponding GENCODE annotations for human and mouse genomes. The resulting gene-level count tables were imported into R version 4.3.3 for downstream analyses. RNA-seq processing metrics are reported in Supplementary Table 6.

### Principal component analysis

PCA was performed to explore global transcriptional differences between groups. PCA was computed using normalized expression data and visualized using standard R packages for multivariate analysis and data visualization.

### Differential expression and functional enrichment analyses

Differential gene expression analyses between groups were performed using DESeq2, which models count data with a negative binomial distribution. Genes were ranked according to |log_2_ Fold-Change>1 and false discovery rate (FDR)-adjusted p values lower than 0.05 to identify significantly up-or down-regulated genes. Gene Set Enrichment Analysis (GSEA) and Gene Ontology (GO) were used to identify pathways and gene sets enriched. Enrichment results were considered significant based on FDR p values lower than 0.25. To estimate pathway activity at the individual sample level, Gene Set Variation Analysis (GSVA) was performed using normalized RNA expression data and Kyoto Encyclopedia of Genes and Genomes (KEGG)^31^ and Hallmark^32^ collections. GSVA scores were calculated for EMT and other relevant signaling pathways. Associations between continuous K_2_ values and baseline GSVA pathway scores were assessed across CDX models using two-sided Spearman rank correlation tests. P values were adjusted for multiple comparisons using the Benjamini-Hochberg procedure, and both nominal P values and false discovery rate-adjusted P values were reported.

### Pathway scoring and machine learning-based predictive modeling

Pathway activity scores were calculated for significantly enriched gene sets from the KEGG^31^ and Hallmark^32^ collections. For each CDX, pathway enrichment scores were computed using GSVA. The top 25 significantly enriched pathway GSVA scores were used as predictors in a LASSO regression model to estimate the K_2_ parameter. Restricting the analysis to this reduced set of pathways, together with LASSO regularization (alpha = 1; gaussian family), was intended to limit model complexity and mitigate overfitting given the small sample size. Model fitting was performed using the glmnet package in R. The optimal regularization parameter (λ) was determined using 10-fold cross-validation to minimize mean squared error. To obtain a more parsimonious and stable model, the value λ.1se (the largest λ within one standard error of the minimum cross-validated error) was selected. A fixed random seed was used to ensure reproducibility. The model was trained using a dataset of 12 CDX models (22RV1, CAPAN-1, ES-2, HCC1937, Hs-746-T, MDA-MB-436, MIA-PaCa-2, NCI-H23, NCI-H460, NCI-H82, SK-OV-3, VCaP). LASSO model inputs and selected coefficients are provided in Supplementary Table 7. K_2_ prediction values are provided in Supplementary Table 8.

### LASSO robustness analyses

LASSO robustness analyses. To assess feature stability, bootstrap stability selection was performed by repeatedly resampling CDX models with replacement and refitting the complete LASSO pipeline. Selection frequency was defined as the proportion of bootstrap iterations in which a pathway retained a non-zero coefficient. Permutation testing was performed by randomly permuting K_2_ labels across CDX models, refitting the LASSO model and comparing the observed model error with the null distribution obtained from permuted datasets. Empirical P values were calculated as the proportion of permuted models showing equivalent or better performance than the observed model. Leave-one-model-out analysis was performed by iteratively excluding a single CDX model, recalculating pathway activity scores for the preselected K_2_-associated pathways. Model performance was then evaluated on the excluded CDX sample. LASSO robustness outputs are reported in Supplementary Table 7 and Extended Data Fig. 1.

### Cross-dataset transfer to CCLE-profiled cell lines

The exploratory cross-dataset transferability of the CDX-trained pathway model was evaluated in an independent set of five cancer cell lines, A2780, HGC-27, IGROV-1, MDA-MB-231, and NCI-H660, for which experimentally determined K_2_ values and transcriptomic profiles were available. GSVA pathway activity scores were calculated from the Cancer Cell Line Encyclopedia (CCLE) RNA-seq profiles using the same gene sets and preprocessing strategy applied to the CDX training dataset. Predicted K_2_ values were generated by applying the coefficients derived from the CDX-trained LASSO model to the corresponding CCLE pathway scores. Model performance was evaluated by comparing predicted and experimentally determined K_2_ values using correlation analysis and normalized mean absolute error (NMAE). Experimental and PHENIX molecular-state score values for the transfer set are provided in Supplementary Table 9.

### Cross-dataset projection of the PHENIX K_2_ model

Publicly available RNA-seq datasets from The Cancer Genome Atlas (TCGA) and the CCLE were used for external projection of the PHENIX pathway model. Following dataset-specific preprocessing, pathway activity scores were calculated using the same gene sets employed during model development. The coefficients derived from the CDX-trained model were then applied to the pathway scores from each dataset to generate predicted K_2_ values for TCGA tumor samples and CCLE cancer cell lines. These predicted values were subsequently used to characterize the distribution of K_2_-associated molecular states across tumor types and cancer cell lineages and to perform the corresponding downstream analyses in each dataset. CCLE and TCGA predicted K_2_ values and projection outputs are reported in Supplementary Tables 11 and 12, respectively.

### Survival and prognostic analyses

Survival analyses were performed in the TCGA cohort using overall survival (OS), event-free survival (EFS), and metastasis-free survival (MFS) as clinical endpoints. Patients were ranked according to their PHENIX-predicted K_2_ values and divided into quartiles. Cases in the first quartile (Q1), corresponding to the highest predicted K_2_ values, were classified as the high-K_2_ group, whereas cases in the fourth quartile (Q4), corresponding to the lowest predicted K_2_ values, were classified as the low-K_2_ group. Patients in the two central quartiles (Q2 and Q3) were combined into a single intermediate-K_2_ group. Kaplan-Meier curves were generated for the three predicted K_2_ groups, and differences in survival distributions were evaluated using log-rank tests, including pairwise comparisons between groups. Univariable Cox proportional hazards models were initially fitted to estimate the association between predicted K_2_ group and each survival endpoint. Multivariable Cox models were subsequently fitted with progressive adjustment for tumor type, pathological stage, and pathological M status. Tumor type was incorporated as a stratification variable to allow the baseline hazard to vary across TCGA cancer types, while estimating a common effect of the predicted K_2_ group. Hazard ratios and corresponding 95% confidence intervals were calculated using the low-K_2_ group as the reference category. The proportional hazards assumption was evaluated using Schoenfeld residuals. Follow-up was censored at six years, and all analyses were performed in R version 4.3.3. All statistical tests were two-sided, and P values ≤ 0.05 were considered statistically significant.

### PDOX transcriptomic pharmacodynamic analysis

For PDOX pharmacodynamic analyses, RNA-seq was performed on tumors collected before treatment (n=3 independent mice), following either vehicle (n=5) or PM534 treatment (n=5). Differential gene expression was nalyzed for two comparisons: vehicle-treated versus pretreatment tumors and PM534-treated versus vehicle-treated tumors. For each comparison, genes were ranked according to their differential expression statistics, and gene set enrichment analysis (GSEA) was performed using the resulting ranked gene lists. Differential expression results are provided in Supplementary Table 13.

## Data availability

The RNA-sequencing data generated in this study have been deposited in the NCBI Sequence Read Archive under BioProject accession PRJNA1541097. All other data supporting the findings of this study are available within the Article and its Supplementary Information or from the corresponding author upon reasonable request.

## Code availability

The custom code used for transcriptomic and pathway analyses, machine learning, cross-dataset projection, survival analyses and figure generation will be deposited in a public repository before publication. Repository details and a permanent identifier will be provided during peer review.

## Author contributions

MR contributed to investigation, methodology, formal analysis and visualization. JR and IF-M contributed to data curation, methodology, software, formal analysis, visualization, and writing, review and editing. RA contributed to investigation, methodology, validation, data curation and resources. MJG contributed to investigation, methodology and resources. A Vidal contributed to investigation, methodology, validation and resources. A Villanueva contributed to investigation, methodology, resources and supervision. CC contributed to conceptualization, supervision, and writing, review and editing. MLR contributed to conceptualization, methodology, formal analysis, supervision, visualization, writing the original draft, and writing, review and editing. PA contributed to conceptualization, investigation, methodology, supervision, project administration, writing the original draft, and writing, review and editing. IFT contributed to conceptualization, methodology, formal analysis, supervision, writing the original draft, and writing, review and editing. All authors reviewed and approved the final version of the manuscript.

## EXTENDED DATA FIGURE LEGENDS

**Extended Data Fig. 1 Robustness analyses of the pathway-based predictor of PM534 sensitivity.** (a) Bootstrap stability analysis showing pathway selection frequencies across resampling iterations. Selection frequency was defined as the proportion of bootstrap iterations in which a pathway retained a non-zero coefficient in the LASSO model. (b) Leave-one-model-out analysis. Predicted, experimental K_2_ values and the corresponding Pearson’s correlation coefficient (r) and normalized mean absolute error (NMAE) are indicated. (c) Permutation analysis of model performance. The observed model MSE and the mean permuted MSE are indicated. (d) Residual diagnostic plot showing residuals (experimental minus predicted K_2_) as a function of predicted K_2_ values for the final pathway-based model.

## SUPPLEMENTARY INFORMATION (as a separate file)

**Supplementary Material 1 -**Additional PBPK–PK/PD model structure, equations and simulation details

**Supplementary Table 1 -** CDX experimental conditions and treatment schedules

**Supplementary Table 2 -** Bioanalytical LC-MS/MS method for PM534 quantification

**Supplementary Table 3 -** PBPK model physiological and estimated parameters

**Supplementary Table 4 -** PK-PD model estimates per each cell line

**Supplementary Table 5.** Endpoint metrics used in CDX models to evaluate K2 against conventional antitumor response phenotypes.

**Supplementary Table 6.** RNA-seq derived molecular metrics across CDX tumor models.

**Supplementary Table 7.** Gene Set Enrichment Analysis results from differential expression analysis of high-k2 CDX tumors compared to low-k2 CDX tumors.

**Supplementary Table 8.** Pathways associated with PM534 pharmacodynamic sensitivity and LASSO feature selection.

**Supplementary Table 9.** Experimental and predicted K_2_ values for training and independent validation cohorts.

**Supplementary Table 10.** Bootstrap stability analysis of candidate pathway predictors.

**Supplementary Table 11.** Model-predicted K_2_ values for CCLE cell lines.

**Supplementary Table 12.** Model-predicted K_2_ values for TCGA tumor samples.

**Supplementary Table 13.** Differential expression analysis of vehicle PDOX tumor compared to pretreatment PDOX tumor (Vehicle / Pretreatment) and PM534 treated PDOX tumor compared to vehicle PDOX tumor (PM534 / Vehicle).

