## Extended Figure 1 for "Pharmacodynamic encoding of tumor exposure identifies a mesenchymal state associated with antitumor sensitivity"

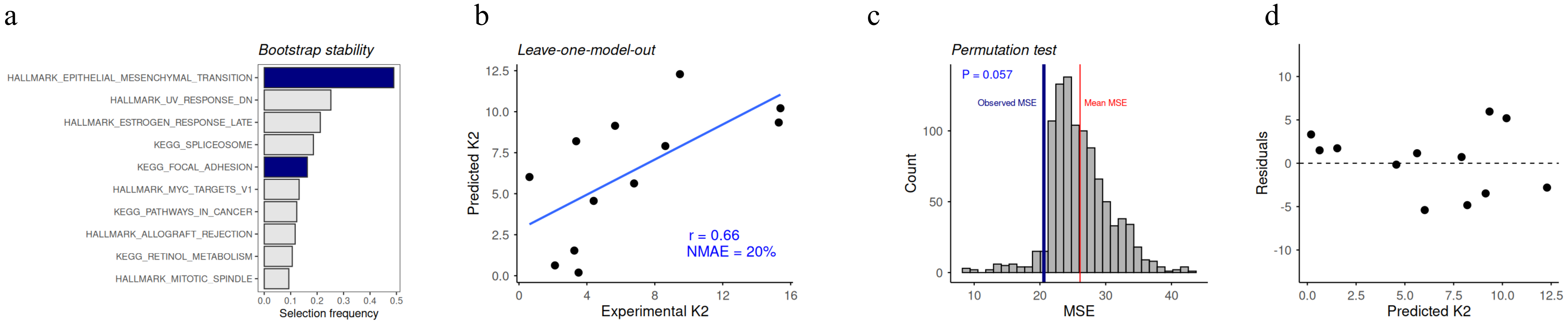

a
Bootstrap stability
HALLMARK_EPITHELIAL_MESENCHYMAL_TRANSITION
HALLMARK_UV_RESPONSE_DN
HALLMARK_ESTROGEN_RESPONSE_LATE
KEGG_SPLICEOSOME
KEGG_FOCAL_ADHESION
HALLMARK_MYC_TARGETS_V1
KEGG_PATHWAYS_IN_CANCER
HALLMARK_ALLOGRAFT_REJECTION
KEGG_RETINOL_METABOLISM
HALLMARK_MITOTIC_SPINDLE
0.0 0.1 0.2 0.3 0.4 0.5
Selection frequency
b
Leave-one-model-out
Predicted K2
Experimental K2
r = 0.66
NMAE = 20%
c
Permutation test
P = 0.057
Observed MSE
Mean MSE
Count
MSE
d
Residuals
Predicted K2
