## Supplementary Material 1 for "Pharmacodynamic encoding of tumor exposure identifies a mesenchymal state associated with antitumor sensitivity"

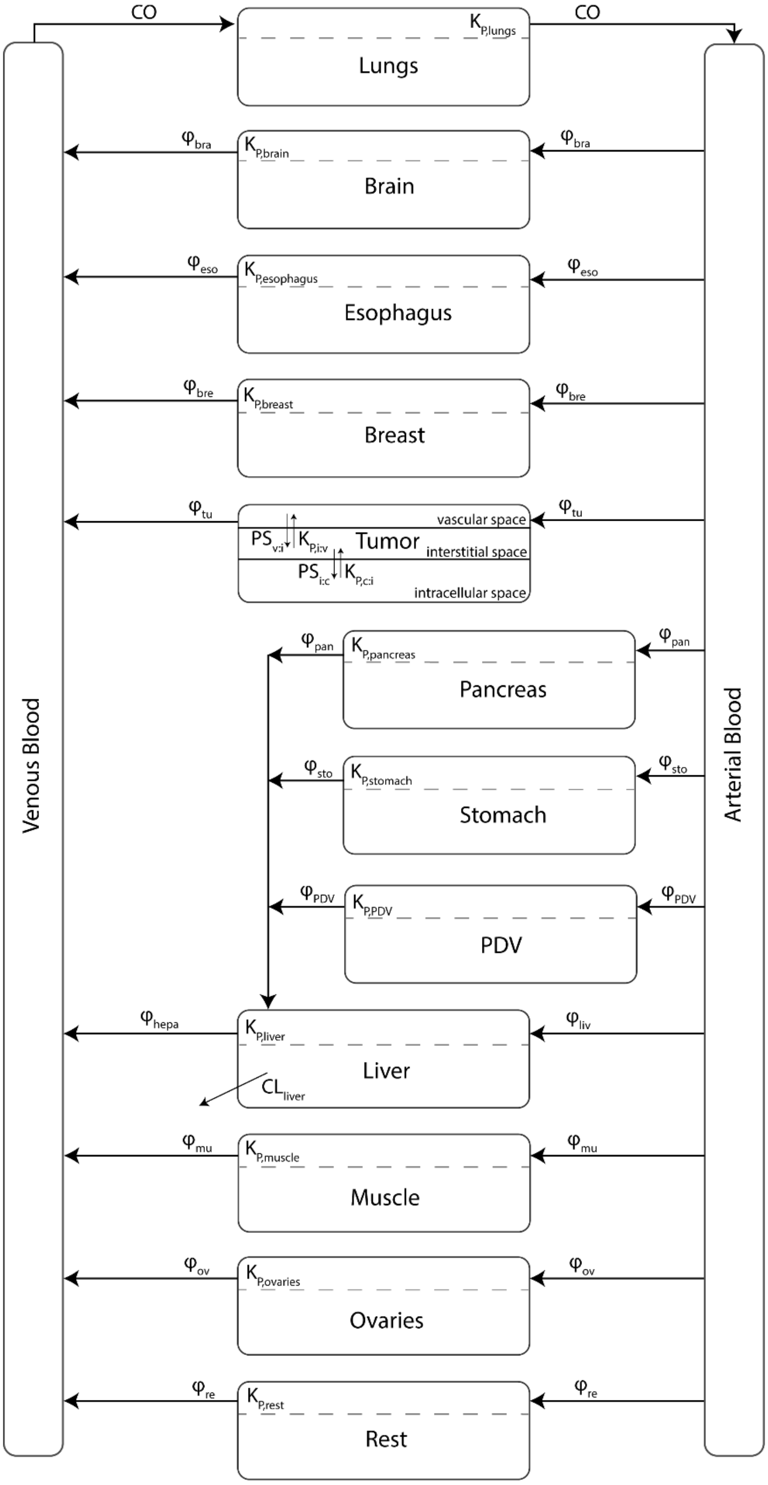


*Figure 1. Physiologically based pharmacokinetic (PBPK) model structure*

**Supplementary Material 1, Figure 1. General PBPK model scheme.** Schematic representation of the physiological based pharmacokinetic model; *CO,* cardiac output; $\varphi$, blood flows; $K_P$, partition coefficients; *PDV*, portal vein-drained visceras; *CL*, clearance; *Rest*, remaining organs (skin, skeleton, heart, etc).

### General tissue expression

$$\frac{dA_{\text{organ}}}{dt} = \varphi_{\text{organ}} \times \left(C_{\text{artery}} - \frac{C_{\text{organ}}}{K_{P,\text{organ}}}\right)$$

### Specific expressions

1. **Veins**

$$\frac{dA_{\text{vein}}}{dt} = \sum\left(\varphi_{\text{organ}} \times \frac{C_{\text{organ}}}{K_{P,\text{organ}}}\right) + \varphi_{\text{tumor}} \times C_{\text{tumor,vas}} - CO \times C_{\text{vein}}$$

2. **Arteries**

$$\frac{dA_{\text{artery}}}{dt} = CO \times \left(\frac{C_{\text{lung}}}{K_{P,\text{lung}}} - C_{\text{artery}}\right)$$

3. **Lungs**

$$\frac{dA_{\text{lung}}}{dt} = CO \times \left(C_{\text{vein}} - \frac{C_{\text{lung}}}{K_{P,\text{lung}}}\right)$$

4. **Liver**

$$\frac{dA_{\text{liver}}}{dt} = IN_{\text{liver}} - (\varphi_{\text{liver}} + \varphi_{\text{stomach}} + \varphi_{\text{pancreas}} + \varphi_{\text{PDV}}) \times \frac{C_{\text{liver}}}{K_{P,\text{liver}}} - CL_{\text{liver}} \times \frac{C_{\text{liver}}}{K_{P,\text{liver}}}$$

$$IN_{\text{liver}} = \varphi_{\text{liver}} \times C_{\text{artery}} + \left(\varphi_{\text{stomach}} \times \frac{C_{\text{stomach}}}{K_{P,\text{stomach}}}\right) + \left(\varphi_{\text{pancreas}} \times \frac{C_{\text{pancreas}}}{K_{P,\text{pancreas}}}\right) + \left(\varphi_{\text{PDV}} \times \frac{C_{\text{PDV}}}{K_{P,\text{PDV}}}\right)$$

5. **Tumor**

**5.1. Vascular**

$$\frac{dA_{\text{tumor,vas}}}{dt} = \varphi_{\text{tumor}} \times (C_{\text{artery}} - C_{\text{tumor,vas}}) - PS_{\text{tumor,iv}} \times C_{\text{tumor,vas}} + PS_{\text{tumor,iv}} \times \frac{C_{\text{tumor,inters}}}{K_{P,\text{iv}}}$$

**5.2. Interstitial**

$$\frac{dA_{\text{tumor,inters}}}{dt} = PS_{\text{tumor,iv}} \times C_{\text{tumor,vas}} - PS_{\text{tumor,iv}} \times \frac{C_{\text{tumor,inters}}}{K_{P,\text{iv}}} - PS_{\text{tumor,ci}} \times C_{\text{tumor,inters}} + PS_{\text{tumor,ci}} \times \frac{C_{\text{tumor,cell}}}{K_{P,\text{ci}}}$$

**5.3. Cellular**

$$\frac{dA_{\text{tumor,inters}}}{dt} = PS_{\text{tumor,ci}} \times C_{\text{tumor,inters}} - PS_{\text{tumor,ci}} \times \frac{C_{\text{tumor,cell}}}{K_{P,\text{ci}}}$$

**Supplementary Material 1, Equations 1. Physiologically based pharmacokinetic (PBPK) model equations**. $A_{\text{x}}$ is the amount of drug in the tissue at a given time; $\varphi_{\text{x}}$ is the blood flow to the tissue; $C_{\text{x}}$ is the drug concentration in the tissue at a given time; $K_{P,\text{x}}$ is the tissue partition coefficient; $CO$ is the cardiac output; $C_{\text{tumor,vas}}$, $C_{\text{tumor,inters}}$ and $C_{\text{tumor,cell}}$ are the drug concentrations in the vascular, interstitial and cellular fractions of the tumor, respectively; $K_{P,\text{iv}}$, $K_{P,\text{ci}}$ are the partition coefficients between vascular and interstitial fractions and interstitial and cellular fractions of the tumor respectively; and $PS_{\text{tumor,iv}}$, $PS_{\text{tumor,ci}}$ are the permeability–surface area products between the vascular and interstitial fractions and between the interstitial and cellular fractions, respectively.
